# Three-dose vaccination-induced immune responses protect against SARS-CoV-2 Omicron BA.2

**DOI:** 10.1101/2022.05.09.491254

**Authors:** Runhong Zhou, Na Liu, Xin Li, Qiaoli Peng, Cheuk-Kwan Yiu, Haode Huang, Dawei Yang, Zhenglong Du, Hau-Yee Kwok, Ka-Kit Au, Jian-Piao Cai, Ivan Fan-Ngai Hung, Kelvin Kai-Wang To, Xiaoning Xu, Kwok-Yung Yuen, Zhiwei Chen

## Abstract

**Background:** The ongoing outbreak of SARS-CoV-2 Omicron BA.2 infections in Hong Kong, the model city of universal masking of the world, has resulted in a major public health crisis. Although the third vaccination resulted in strong boosting of neutralization antibody, vaccine efficacy and corelates of immune protection against the major circulating Omicron BA.2 remains to be investigated.

**Methods:** We investigated the vaccine efficacy against the Omicron BA.2 breakthrough infection among 470 public servants who had received different SARS-CoV-2 vaccine regimens including two-dose BNT162b2 (2×BNT, n=169), three-dose BNT162b2 (3×BNT, n=170), two-dose CoronaVac (2×CorV, n=34), three-dose CoronaVac (3×CorV, n=67) and third-dose BNT162b2 following 2×CorV (2×CorV+1BNT, n=32). Humoral and cellular immune responses after three-dose vaccination were further characterized and correlated with clinical characteristics of BA.2 infection.

**Findings:** During the BA.2 outbreak, 27.7% vaccinees were infected. The timely third-dose vaccination provided significant protection with lower incidence rates of breakthrough infections (2×BNT 49.2% vs 3×BNT 13.1%, p <0.0001; 2×CorV 44.1% vs 3×CoV 19.4%, p=0.003). Investigation of immune response on blood samples derived from 92 subjects in three-dose vaccination cohorts collected before the BA.2 outbreak revealed that the third-dose vaccination activated spike (S)-specific memory B cells and Omicron cross-reactive T cell responses, which correlated with reduced frequencies of breakthrough infections and disease severity rather than with types of vaccines. Moreover, the frequency of S-specific activated memory B cells was significantly lower in infected vaccinees than uninfected vaccinees before vaccine-breakthrough infection whereas IFN-γ^+^ CD4 T cells were negatively associated with age and viral clearance time. Critically, BA.2 breakthrough infection boosted cross-reactive memory B cells with enhanced cross-neutralizing antibodies to Omicron sublineages, including BA.2.12.1 and BA.4/5, in all vaccinees tested.

**Interpretation:** Our results imply that the timely third vaccination and immune responses are likely required for vaccine-mediated protection against Omicron BA.2 pandemic. Although BA.2 conferred the highest neutralization resistance compared with variants of concern tested before the emergence of BA.2.12.1 and BA.4/5, the third dose vaccination-activated S-specific memory B cells and Omicron cross-reactive T cell responses contributed to reduced frequencies of breakthrough infection and disease severity. Neutralizing antibody potency enhanced by BA. 2 breakthrough infection with previous 3 doses of vaccines (CoronaVac or BNT162b2) may reduce the risk for infection of ongoing BA.2.12.1 and BA.4/5.

**Funding:** Hong Kong Research Grants Council Collaborative Research Fund, Health and Medical Research Fund, Wellcome Trust, Shenzhen Science and Technology Program, the Health@InnoHK, Innovation and Technology Commission of Hong Kong, China, National Program on Key Research Project, Emergency Key Program of Guangzhou Laboratory, donations from the Friends of Hope Education Fund and the Hong Kong Theme-Based Research Scheme.

## Introduction

To fight the ongoing SARS-CoV-2 pandemic, over 10 billion doses of COVID-19 vaccines under emergency use authorization (EUA) have been administered globally, which has significantly reduced the rates of hospitalization, disease severity and death ^1–5^. Unfortunately, the emergence of variants of concern (VOCs), especially the Omicron variants, have substantially threatened the vaccine efficacy ^6^. We recently reported that waning anti-Omicron neutralizing antibody and T cell responses especially among CoronaVac-vaccinees might pose a risk to vaccine-breakthrough infections in Hong Kong ^7^. Although the third heterologous BNT162b2 vaccination after 2-dose CoronaVac generates high neutralizing antibody responses against ancestral and Omicron BA.1 than the third homologous CoronaVac booster ^8,9^, vaccine efficacy and its correlations with the immune protection against the major circulating Omicron BA.2 remains to be investigated ^10–12^ In addition, it remains unclear if BA.2 breakthrough infection would reduce the risk against ongoing BA.2.12.1 and BA.4/5 reinfection by enhancing cross-reactive neutralizing antibody potency.

## Materials and methods

### Human subjects

This study was approved by the Institutional Review Board of the University of Hong Kong/Hospital Authority Hong Kong West Cluster (Ref No. UW 21-452). A total of 481 participants were recruited in this study. Written informed consent and questionnaire of vaccination and infection were obtained from these subjects. Patients provided the information of symptom onset date, type of symptoms, hospitalization, duration of illness and the date of viral negative conversion as summarized in Table 1. The vaccination record was officially registered by professional medical staff in the governmental system called “LeaveHomeSafe”. The diagnosis of SARS-CoV-2 infection was confirmed by results of rapid antigen test and PCR, as well as quarantine records enforced strictly by law. Peripheral blood mononuclear cells (PBMCs) from 92 randomly selected-participants who had the third vaccination were isolated from fresh blood samples before SARS-CoV-2 infection using Ficoll-Paque density gradient centrifugation in our BSL-3 laboratory at the same day of blood collection. The majority of purified PBMCs were used for immune cell phenotyping whereas plasma samples were subjected to antibody testing. The rest of the cells were cryopreserved in freezing medium (Synth-a-Freeze Cryopreservation Medium, ThermoFisher Scientific) at 5×10^6^ cells/mL at −150°C. Subjects included in the study were required to complete vaccination (all dose) for at least 7 days, to allow the manifestation of the delayed immune response to vaccination.

**Table 1.** Demographic characteristics of breakthrough infection among 470 vaccinees

| Vaccinations |  | 2×BNT<br>(n=169) | 3×BNT<br>(n=168) | 2×CorV<br>(n=34) | 3×CorV<br>(n=67) | 2×CorV+1×BNT<br>(n=32) |
| --- | --- | --- | --- | --- | --- | --- |
| Infection rate %<br>(No. patient/Total No.) |  | 49.2%<br>(78/169) | 13.1%<br>(22/168) | 44.1%<br>(15/34) | 19.4%<br>(13/67) | 6.3%<br>(2/32) |
| Patients |  | (n=78) | (n=22) | (n=15) | (n=13) | (n=2) |
| Age, year<br>(ranges in parentheses) |  | 32<br>(24-58) | 40<br>(27-60) | 41<br>(24-64) | 50<br>(20-62) | 47.5<br>(37-58) |
| Gender | Male (% of all participants) | 60 (48.8%) | 14 (12.3%) | 9 (42.9%) | 8 (18.2%) | 2 (7.1%) |
|  | Female (% of all participants) | 18 (39.1%) | 8 (14.8%) | 6 (46.2%) | 5 (21.7%) | 0 (0%) |
| Median interval days<br>between latest vaccination<br>and symptom onset (ranges<br>in parentheses) |  | 227<br>(140-332) | 48.5<br>(10-111) | 237<br>(52-341) | 56<br>(7-109) | 25.5<br>(10-41) |
| Asymptomatic rate %<br>(No. Asymptomatic<br>patient/No. total patient) |  | 3.8%<br>(3/78) | 0%<br>(0/22) | 0 %<br>(0/15) | 0%<br>(0/13) | 0%<br>(0/2) |
| Disease severity |  | Mild | Mild | Mild | Mild | Mild |
| Number of symptoms<br>(ranges in parentheses) |  | 4<br>(0-6) | 3<br>(1-5) | 3<br>(1-6) | 2<br>(1-5) | 3.5<br>(3-5) |
| Presentation to hospital %<br>(No. patients presenting to<br>hospital/No. total patient) |  | 19.2%<br>(15/78) | 4.5%<br>(1/22) | 20%<br>(3/15) | 15.4%<br>(2/13) | 50%<br>(1/2) |
| Duration of illness, days<br>(ranges in parentheses) |  | 7<br>(0-19) | 7.5<br>(2-19) | 8<br>(6-21) | 8<br>(2-14) | 9.5<br>(2-17) |
| The interval days between<br>symptom onset and two<br>negative RAT |  | 8<br>(1-20) | 9<br>(6-13) | 8<br>(6-12) | 9<br>(3-14) | 8<br>(5-11) |
| Values displayed are medians, with ranges in parentheses |  |  |  |  |  |  |

### Enzyme-linked immunosorbent assays (ELISA)

Serum IgG binding antibodies to Spike were quantitated by ELISA using WHO International Standard as standard. Briefly, different recombinant trimeric Spike proteins derived from SARS-CoV-2 VOCs (Sino Biological) were diluted to final concentrations of 1 μg/mL, then coated onto 96-well plates (Corning 3690) and incubated at 4 °C overnight. Plates were washed with PBST (PBS containing 0.05% Tween-20) and blocked with blocking buffer (PBS containing 5% skim milk or 1% BSA) at 37 °C for 1 h. Two-fold serial dilution of WHO international standard (from 20 BAU/mL to 0.15625 BAU/mL) and plasma samples (400-fold diluted) were added to the plates and incubated at 37 °C for 1 h. Wells were then incubated with a secondary goat anti-human IgG labeled with horseradish peroxidase (HRP) (1:5000 Invitrogen) TMB substrate (SIGMA). Optical density (OD) at 450 nm was measured by SkanIt RE6.1 with VARIOSKAN Lux (Thermo Scientific).

### Pseudotyped viral neutralization assay

To determine the neutralizing activity of subject’s plasma, the plasma was inactivated at 56°C for 30 min prior to a pseudotyped viral entry assay. In brief, different SARS-CoV-2 pseudotyped viruses were generated through co-transfection of 293T cells with 2 plasmids, pSARS-CoV-2 S and pNL4-3Luc_Env_Vpr, carrying the optimized SARS-CoV-2 S gene and a human immunodeficiency virus type 1 backbone, respectively. At 48 h post-transfection, viral supernatant was collected and frozen at −150°C. Serially diluted plasma samples (from 1:20 to 1:14580) were incubated with 200 TCID50 of pseudovirus at 37°C for 1 h. The plasma-virus mixtures were then added into pre-seeded HEK293T-hACE2 cells. After 48 h, infected cells were lysed, and luciferase activity was measured using Luciferase Assay System kits (Promega) in a Victor3-1420 Multilabel Counter (PerkinElmer). The 50%inhibitory concentrations (IC_50_) of each plasma specimen were calculated to reflect anti-SARS-CoV-2 potency.

### Antigen-specific B cells

To characterize the SARS-CoV-2 Spike-specific B cells, PBMCs from each vaccinee were first stained with an antibody cocktail contained dead cell dye (Zombie aquae), CD3-Pacific Blue, CD14-Pacific Blue, CD56-Pacific Blue, CD19-BV785, IgD-BV605,IgG-PE, CD27-BV711, CD21-PE/Cy7, CD38-Percp/Cy5.5, CD11C-APC/Fire750 and His-tag Spike protein. Cells were then washed with FACS buffer (PBS with 2% FBS) and further stained with the secondary antibodies including APC anti-His and DyLight 488 anti-his antibodies. Stained cells were acquired by FACSAriaIII Flow Cytometer (BD Biosciences) and analyzed with FlowJo software (v10.6) (BD Bioscience).

### Intracellular cytokine staining (ICS)

To measure antigen-specific T cell responses, PBMCs were stimulated with 2 μg/mL Spike peptide pool (15-mer overlapping by 11) from SARS-CoV-2 ancestral or Omicron variant, or 2 μg/mL nucleocapsid protein (NP) peptide pool in the presence of 0.5 μg/mL anti-CD28 and anti-CD49d mAbs (Biolegend). Cells were incubated at 37°C for 9 hours and BFA was added at 3 h post incubation, as previously described ^11^. PMA/ionomycin stimulation was included as positive control. Cells were then washed with staining buffer (PBS containing 2% FBS) and stained with mAbs against surface markers, including dead cell dye (Zombie aqua), CD3-Pacific Blue, CD4-Percp/Cy5.5, CD8-APC/Fire750, CD45RA-BV711, CCR7-BV785, CXCR5-APC, CCR6-BV605. For intracellular staining, cells were fixed and permeabilized with BD Cytofix/Cytoperm (BD Biosciences) prior to staining with the mAbs against IFN-γ-PE, TNF-α-AF488 and IL-2-PE-Cy7. Stained cells were acquired by FACSAriaIII Flow Cytometer (BD Biosciences) and analyzed with FlowJo software (v10.6) (BD Bioscience). Results were subtracted from percentage of unstimulated control.

### Correlation plots and heatmap visualizations

Correlograms plotting the Spearman rank correlation coefficient (r), between all parameter pairs were generated with the corrplot package (v0.84) ^13^ running under R (v3.6.1) in RStudio (1.1.456). Spearman rank two-tailed P values were calculated using corr.test (psych v1.8.12) and graphed (ggplot2 v3.1.1) based on *p<0.05, **p<0.01, ***p<0.001.

### Statistical analysis

Statistical analysis was performed using PRISM 8.0. For between-group or multiple-group categorical values comparison, two-sided chi-square tests or fisher’s exact tests were used. Unpaired Student’s t tests were used to compare group means of GMT and cell frequencies between two groups. The statistic details are depicted in the respective legends. A P value <0.05 was considered significant.

## Results

### Demographic characteristics of breakthrough infection among 481 vaccinees

Considering sociodemographic characteristics and exposure risk may also affect vaccine efficacy. In this study, therefore, we only focus on 7247 subjects who are public servants working for Hong Kong Government with comparable exposure risks. During the time from January to March 2022 (Omicron BA.2 was first found in mid-January 2022 and reached the peak in the early March as dominant strain in Hong Kong^10,14^), 5995 (82.7%) and 1012 (14%) study subjects had received two and three doses of vaccinations, respectively, resulting in an overall vaccination rate of 96.7%. During the recent fifth wave of COVID-19 in Hong Kong since the end of January 2022 ^10^, 470 (6.5%) subjects joined our follow-up study. These subjects had received 2-dose BNT162b2 (2×BNT, n=169), 3-dose BNT162b2 (3×BNT, n=168), 2-dose CoronaVac (2×CorV, n=34), 3-dose CoronaVac (3×CorV, n=67) or a heterologous booster dose of BNT162b2 after two prior doses of CoronaVac (2×CorV+1×BNT, n=32) (Table 1). Among these 470 subjects, a total of 141 (128/470, 27.2%) infections were confirmed by governmental reverse transcriptase-polymerase chain reaction (RT-PCR) or lateral flow-based rapid antigen test (RAT) during the study period. Gender difference in infection was not observed. Patients in 2×BNT were relatively younger than 3×BNT (2×BNT vs 3×BNT: median 32 years vs median 40 years, p<0.0001), likely indicating the hesitation for taking the third dose BNT162b2 among younger people. Patients who received two dose BNT162b2 were significantly younger than patients who received two dose CoronaVac (2×CorV vs 2×BNT: median 41 years vs median 32 years, p=0.0006 (Table 1 and Supplementary Table 1), in line with elderly people’s preference of taking CoronaVac with less side effects. Moreover, a shorter median interval between latest vaccination and symptom onset was noticed for 3×BNT compared to 2×BNT (2×BNT vs 3×BNT: median 227 days vs median 48.5 days, p<0.0001) and for 3×CorV compared to 2×CorV (2×CorV vs 3×CorV: median 237 days vs median 56 days, p<0.0001), respectively (Table 1 and Supplementary Table 1).

Infections were found in both 2×BNT and 2×CorV groups with comparable incidence rates of 49.2% (78/169) and 44.1% (15/34) (*p*=0.828), respectively. For the third dose vaccination groups, however, both third homologous BNT162b2 (3×BNT: 22/168, 13.1%, p<0.0001) and CoronaVac vaccination (3×CorV: 13/67, 19.4%, p=0.009) showed significantly reduced infection rate compared to 2×BNT and 2×CorV, respectively. The third heterologous BNT162b2 vaccination group (2×CorV+1×BNT) exhibited the lowest incident rate of 6.3% compared to the 2×CorV group (*p*<0.0001). No statistical significance was found in the infection rates between any 3 dose groups, although 3×BNT and 2×CorV+1×BNT showed lower infection rates than 3×CorV (Table 1 and Supplementary Table 1). Notably, most infected subjects developed mild disease, presenting three major symptoms including fever, cough and/or sore throat. Asymptotic infections were only found in 2×BNT groups with a low frequencies of 3.8% (3/78) (Table 1). The hospitalization rate was lower for 3×BNT (4.5%) than that of 3×CorV (15.4%) patients. Comparable illness duration was observed in 2×BNT (median 7 days) and 3×BNT (median 7.5 days) than those of 2×CorV (median 8 days) and 3×CorV (median 8 days). There was no significant difference in terms of duration time for viral antigen conversion to negativity between any groups (Table 1 and Supplementary Table 1). These results suggested that the third dose vaccination by both BNT162b2 and CoronaVac reduced the incident rate of BA.2 infection and the third dose of BNT162b2 vaccination achieved a slightly lower hospitalization rate compared with the third CoronaVac.

### Activation of Spike-specific memory B cells by the third vaccination

To characterize the third dose vaccination-induced immune responses, we were able to obtain 92 blood samples donated by subjects in the same cohort including 41 from 3×BNT, 28 from 3×CorV and 21 from 2×CorV+1×BNT at median 23, 56 and 47 days after the last vaccination, respectively, on January 27, 2022, right before BA.2 outbreak in Hong Kong ^10,14^(Supplementary Table 2). Considering that memory B cell responses contribute to long-term immunological protection against COVID-19, we measured the frequency of Spike (S)-specific B cells (gated on CD19^+^ IgG^+^ IgD^-^ cells) after the third dose vaccination (Figure 1A). We found that the third dose of BNT162b2, either 3×BNT (mean 2.83%) or 2×CorV+1×BNT (mean 1.33%), induced significant higher frequency of S-specific B cells than 3×CorV (mean 0.35%) (Figure 1B). The significant boost effect of S-specific B cells was not observed by the third dose of CoronaVac (Figure 1C). Moreover, S-specific B cells elicited by the third dose of BNT162b2 reached the peak around 4-6 weeks and lasted for 3 months with a higher mean frequency than that of 3×CorV (Figure 1D). Further phenotypical analysis (Figure 1E) showed that the third dose of BNT162b2 resulted in elevated frequency of activated memory B cells (AM, CD21^-^CD27^+^) compared with 2×BNT or 2×CorV whereas the third dose of CoronaVac enhanced the frequency of resting memory (RM) B cells (Figure 1F). The frequency of AM reached the peak at 4 weeks after the third booster and subsequently declined, accompanied by proportional increase of RM, in both 3×BNT and 2×CorV+1×BNT groups whereas AM remained unchanged in the 3×CorV group around two months (Figure 1G). These results demonstrated that S-specific memory B cells were predominantly activated by the third dose of BNT162b2 but insignificantly by the third dose of CoronaVac. However, the third BNT162b2 vaccination following 2 doses of CoronaVac-boosted S-specific B cells was comparable to those induced by three doses of BNT162b2, indicating that BNT162b2 can recall and augment CoronaVac-induced memory B cells.

**Figure 1.**
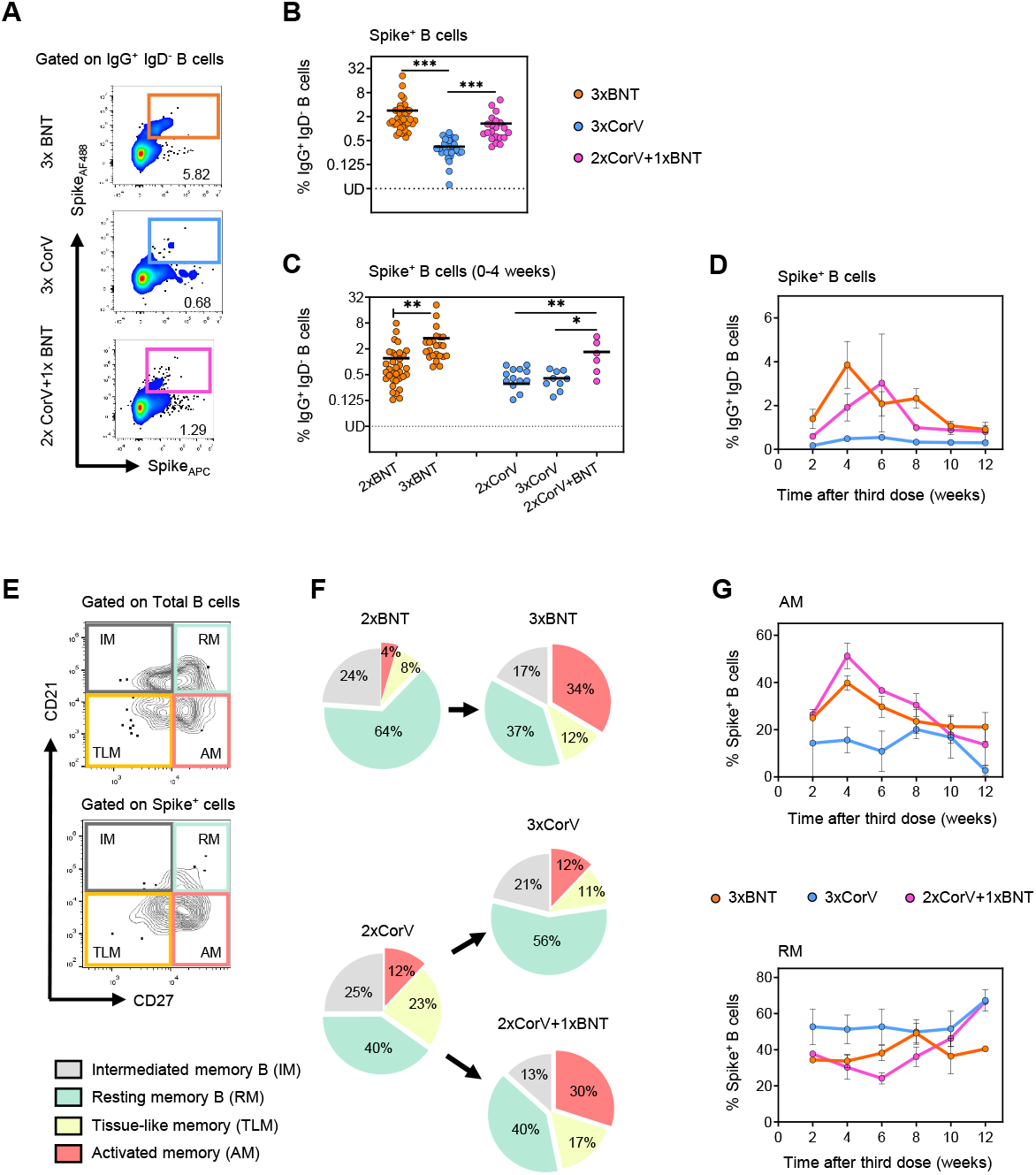
Activation of Spike-specific memory B cells by the third dose vaccination. (**A**) Representative flow cytometry plots showing staining patterns of SARS-CoV-2 Spike probes on memory B cells (IgD^-^ IgG^+^ CD19^+^). (**B**) Quantified results depict the percentage of Spike^+^ B cells in 3×BNT (orange), 3×CorV (blue) and 2×CorV+1×BNT (purple) groups at median 23, 55 and 47 days after the third dose vaccination. (**C**) Comparisons of Spike^+^ B cell frequency between 2-dose (sample collected at median 28 days after second vaccination for 2×BNT and 2×CorV groups) and 3-dose (sample collected at median 16, 20 and 18.5 days after third vaccination for 3×BNT, 3×CorV and 2×CorV+1×BNT groups, respectively) cohorts within 4 weeks after the last vaccinations. (**D**) Cross-sectional analysis of Spikespecific B cells by time after third dose vaccination. The connection lines indicate the mean value. (**E**) Phenotypes of Spike-specific B cells were defined by using CD21 and CD27 markers. (**F**) Pie chart showed the proportion of activated (AM), tissue like (TLM) memory, intermediate memory (IM) and resting-memory (RM) B cells. (**G**) Cross-sectional analysis of the percentage of AM (upper) and RM (bottom) in the Spike-specific B cells by time after third vaccination. The connection lines indicate the mean value.

### The titer and breadth of neutralizing antibodies (NAbs) against a full panel of current SARS-CoV-2 VOCs

We then measured the titer and breadth of neutralizing antibodies (NAbs) against a full panel of current SARS-CoV-2 VOCs including D614G, Alpha, Beta, Delta and five Omicron variants (BA.1, BA.1.1, BA.2, BA.2.12.1 and BA.4/5) using the pseudovirus assay as we previously described ^7^. We included data from subjects who previously received 2×BNT or 2×CorV at the activation (0-4 weeks) and memory (4-15 weeks) periods were used for comparison (Supplementary Table 2). In line with significantly higher frequencies of S-specific B cells, both 3×BNT- and 2×CorV+1×BNT-vaccinees displayed significantly stronger geometric mean 50% neutralizing titers (GMT) than 3×CorV against all variants tested (Figure 2A). The overall fold of neutralization resistance followed the order of Alpha < Beta < Delta < Omicron lineages in all three vaccine groups. Interestingly, Omicron BA.2 and BA. 4/5 were more resistant to other VOCs with comparable reduction fold of GMT while BA.2.12.1 showed a downward resistance compared to BA.2 among all vaccinees (Figure 2B). According to the criteria that convalescent plasma antibody titer >1:320 were eligible initially for SARS-CoV-2 therapy ^15^ and considering that the prophylactic administration of convalescent plasma at 1:320 dilution hardly prevents SARS-CoV-2 infection in the hamster model ^16^, we used 1:320 as the threshold to define NAb titer: less than 1:320 as “Low”, 1:320-1:1280 as “Medium” and above 1: 1280 as “High” for proportion analysis (Figure 2C). We found that 61% of 3×BNT and 48% of 2×CorV+1×BNT vaccinees had high neutralization activity (>1280) against D614G whereas none of 3×CorV vaccinees showed similar activities (Figure 2C). For BA.2, neither 3×BNT nor 2×CorV+1×BNT vaccinees had high neutralization activity, but 41% of 3×BNT and 29% of 2×CorV+1×BNT vaccinees still had medium neutralization activity (321-1280). Strikingly, 68% of 3×CorV vaccinees showed undetectable neutralization antibodies against BA.2. Similar proportion of GMT magnitude was observed in all vaccine groups against BA.4/5 (Figure 2C). We also compared the binding antibody titers using different VOC spike protein as the coating antigen. Since spike-specific IgG titers were correlated positively with the neutralizing potency ‘, we found that sera binding titers of various VOCs in 3×BNT and 2×CorV+1×BNT groups were dramatically higher than those in 3×CorV group (Figure 2D). However, as vaccine-induced NAbs wane over time ^7^, we further compared the NAb titer between 2-dose and 3-dose vaccinees at the similar time post-vaccination (without significant difference) (Supplementary Table 3). The third dose of BNT162b2 induced significant higher NAb titers against all VOCs in 3×BNT and 2×CorV+1×BNT groups compared to the 2-dose groups at both 0-4 weeks (activation) and >4 weeks (memory) after vaccination (Supplementary Table 3). In contrast to weak boost effects by the third dose of CoronaVac in the 3×CorV group, 10.1-26.1-fold and 9.7-27.5-fold enhancements against Omicron variants at activation and memory phases were observed after the third heterologous BNT162b2 (2×CorV+1×BNT), similar to the boost effects in the 3×BNT group (Supplementary Table 3). Apart from the significantly increased NAb titers, the responder rates of anti-BA.2 raised from 33% to 100%, from 0% to 38% and from 0% to 100% at 0-4 weeks; from 39% to 100%, from 0% to 35% and from 0% to 100% at >4 weeks in 3×BNT, 3×CorV and 2×CorV+1×BNT groups, respectively, post the last vaccination. Consistently, BA.2 exhibited the most resistant profile to the boost effect, especially in 3×CorV (Supplementary Table 4). These results demonstrated that the third heterologous BNT162b2 vaccination in 2×CorV+1×BNT made significant improvement on not only bringing the anti-Omicron responder rate to 100% but also enhancing NAb titers close to 3×BNT at both 0-4 and >4 weeks (Supplementary Table 3 and Supplementary Table 4).

**Figure 2.**
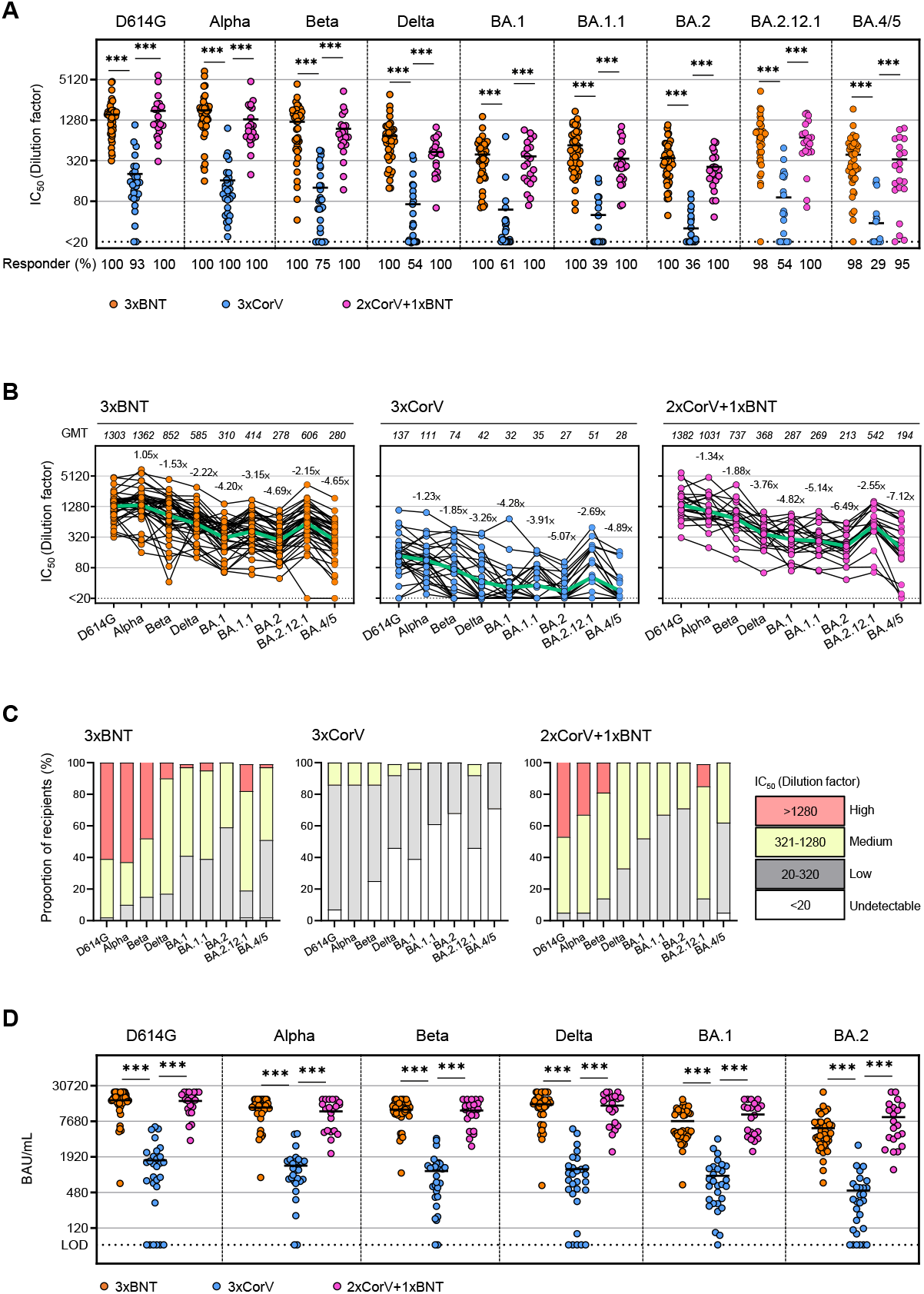
The titer and breadth of neutralizing antibodies (NAbs) against a full panel of current SARS-CoV-2 VOCs. (**A**) The geometric mean titers (GMT) of neutralizing antibody (IC_50_ represents serum dilution required to achieve 50% virus neutralization) against nine SARS-CoV-2 strains were measured by pseudovirusbased assay among 3×BNT (orange), 3×CorV (blue) and 2×CorV+1×BNT vaccinees (purple) at median 23, 55 and 47 days after the third dose vaccination. Numbers under the x-axis indicate the responder rates (IC50>20 termed ‘responder’). (**B**) GMT of neutralizing antibody were depicted on the top of Figure. The green lines indicate the change of GMT among variants. Numbers on the top of dots indicate the fold change of different VOC relative to D614G. Each symbol represents an individual donor with a line indicating the mean of each group. (**C**) Proportion of four neutralizing antibody magnitudes among vaccinees. (**D**) Levels of anti-Spike IgG (BAU/mL) of all vaccinated subjected are shown as mean ± SEM. Dotted line represents value of 64.5 BAU/mL used as the limit of detection (LOD). Statistics were generated by using 2tailed Student’s t test. *p<0.05; **p<0.01; ***p<0.001.

### Spike-specific CD4 and CD8 T cell responses

T cell responses may play an important role in control of SARS-CoV-2 infection ^11,12,17^, CD4 and CD8 T cell responses to viral Spike (S) and nucleocapsid protein (NP) were determined by measuring intracellular IFN-γ, TNF-α and IL-2 (Figure 3A and 3E). Since many amino acid mutations were found in Omicron Spike protein, we measured ancestral and Omicron S-specific T cell responses in parallel. Significantly higher mean frequencies of S-specific IFN-γ^+^ CD4 T cells were found in 3×BNT (ancestral: 0.070% and Omicron: 0.080%) than those in 3×CorV (ancestral: 0.025% and Omicron: 0.023%) and in 2×CorV+1×BNT (ancestral: 0.034% and Omicron: 0.030%) (Figure 3B). No significant differences of S-specific IFN-γ^+^ and polyfunctional CD4 T cells were found between ancestral and Omicron (Figure 3B and 3C). There were also no significant differences between 2×BNT and 3×BNT, and between 2×CorV and 3×CorV at activation period (Figure 3D, left). However, the third BNT162b2 vaccination in the 2×CorV+1×BNT group recalled significant higher frequency of S-specific IFN-γ^+^ cells and responder rate than those in the 3×CorV group at the memory phase (Figure 3D, right). In addition, significantly higher mean frequencies of S-specific IFN-γ^+^ CD8 T cells were found in 3×BNT (ancestral: 0.084% and Omicron: 0.098%) than those in 3×CorV (ancestral: 0.017% and Omicron: 0.015%) and in 2×CorV+1×BNT (ancestral: 0.021% and Omicron: 0.013%) (Figure 3F). The frequency of S-specific polyfunctional CD8 T cells were relatively higher in 3×BNT than those in 3×CorV and 2×CorV+1×BNT (Figure 3G). Significant increase of S-specific IFN-γ^+^ CD8 T cells was not observed in 3×BNT compared to that in 2×BNT at acute (Figure 3H, left) but observed at the memory period (Figure 3H, right). CoronaVac, however, did not show similar activities. Besides the Spike, weak nucleocapsid protein (NP)-specific IFN-γ^+^ CD4 and CD8 T cells were observed in 3 groups although more CD4 T cell responders (67%) were found in 3×CorV (Supplementary Figure 1), indicating the pre-existing of cross-reactive NP-specific T cell responses in unexposed donors ^18^. Considering that S-specific circulating T follicular helper cells (cTFH, CD45RA^-^CXCR5^+^CD4^+^) are associated with potent NAb responses ^19^, we found that the frequency of IFN-γ^+^ cTFH cells were low with mean 0.033-0.048%, 0.01-0.023% and 0.017-0.059% in 3×BNT, 3×CorV and 2×BNT+1×CorV groups, respectively (Supplementary Figure 2A-2B). However, the responder rate was higher in 3×BNT (20-22%) and 2×BNT+1×CorV (14-24%) than that of 3×CorV (7-10%) (Supplementary Figure 2B). These results indicated that the third dose of BNT162b2 vaccination significantly improved S-specific IFN-γ^+^, polyfunctional and memory T cells in 3×BNT but not in 2×CorV+1×BNT and 3×CorV.

**Figure 3.**
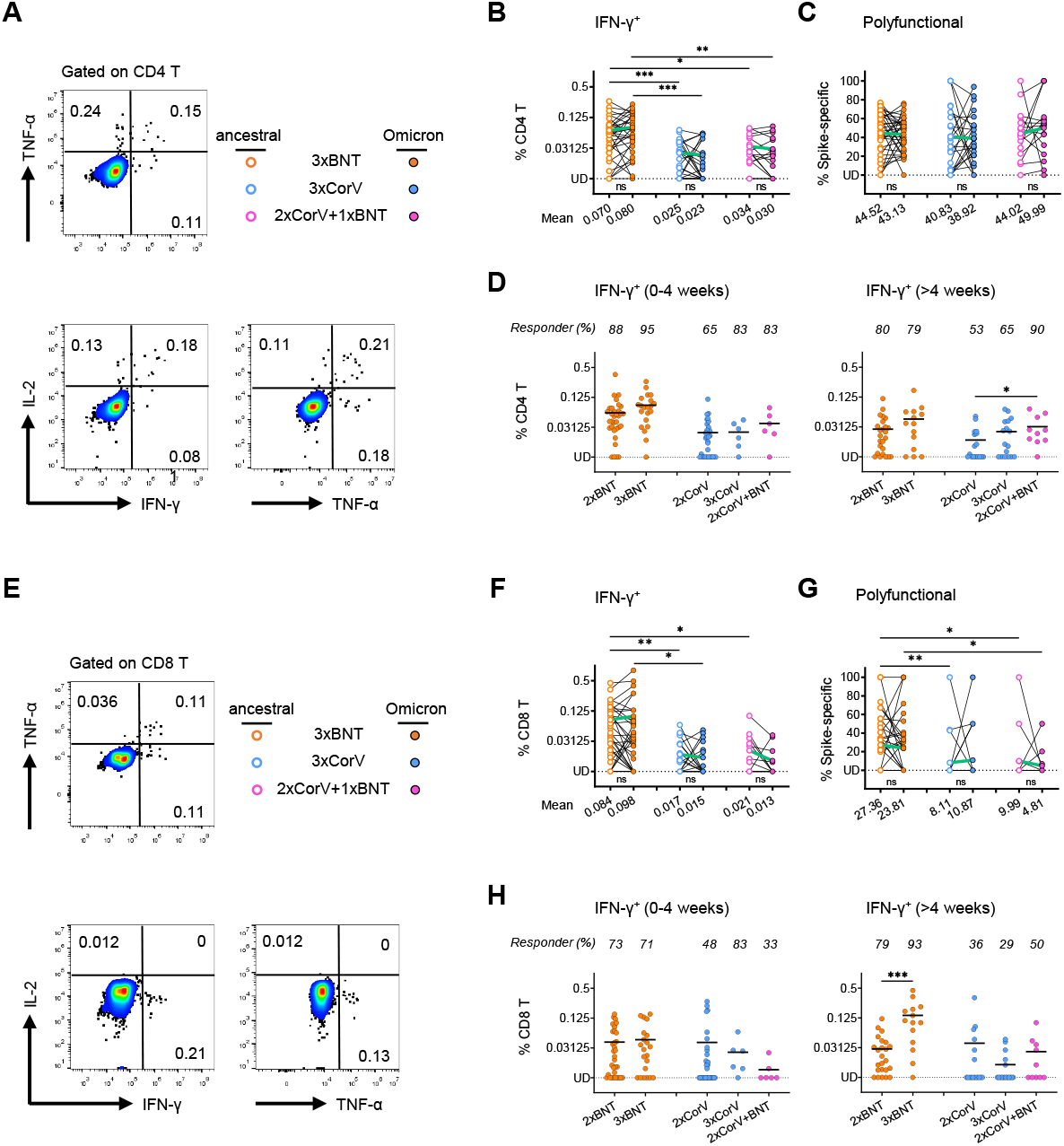
Spike-specific CD4 and CD8 T cell responses. PBMCs were stimulated with the Spike peptide pools from ancestral or Omicron SARS-CoV-2 prior to intracellular cytokine staining assay. Representative flow cytometry plots showing single positive of IFN-γ^+^ or TNF-α^+^ or IL-2^+^ as well as the polyfunctional cells with >2 cytokines among CD4^+^ (**A**) and CD8^+^ (**E**) T cells. Paired analysis of the frequencies of IFN-γ-producing CD4^+^ (**B**) and CD8^+^ (**F**) T cells as well as the frequencies of polyfunctional CD4^+^ (**C**) and CD8^+^ (**G**) T cells to ancestral (open dots) or Omicron (solid dots) Spike among the 3×BNT (orange), 3×CorV (blue) and 2×CorV+1×BNT (purple) vaccinees. The mean frequencies were depicted under the x-axis. The frequencies of IFN-γ-producing CD4^+^ (**D**) and CD8^+^ (**H**) T cell to ancestral Spike among 2×BNT, 3×BNT, 2×CorV, 3×CorV and 2×CorV+1×BNT vaccinees at 04 weeks (left) and >4 weeks (right) periods after vaccinations. Undetected (UD): % of IFN-γ^+^ cells<0.00781%. The green lines in **B**, **C**, **F**, **G**indicate the change of mean responses to ancestral and Omicron Spike. The responses are depicted as the background-subtracted percentage of S-specific T cells (Background subtraction refers to the subtraction of the values of the negative control sample from the peptide-stimulated sample). The responder rates were depicted on the top of dots (% of IFN-γ^+^cells>0.00781% termed ‘responder’ after subtracted from percentage of unstimulated control). Each symbol represents an individual donor with a line indicating the mean of each group. Statistics were generated by using 2-tailed Student’s t test. Ns: no significance, *p<0.05; **p<0.01; ***p<0.001.

### Associations among humoral, cellular immune response and breakthrough infection features

Immune correlation analysis was subsequently conducted for 23 antigen-specific measurements together with gender, age, time interval between second and third vaccinations, sampling time after third dose of vaccination and infection. Consistent with the kinetics of AM proportion, S-specific AM correlated negatively with time after the third dose of vaccination in the 2×CorV+1 ×BNT group (Figure 4C). Positive correlations between S-specific B cells and NAbs were observed in both 3×BNT and 2×CorV+1 ×BNT groups while the RM was positively associated with NAbs in the 3×CorV group (Figure 4A-C, green rectangle). Consistently, significant positive correlations were found in NAbs titers against all 7 viral variants (Figure 4A-C, purple triangles). Since the third dose vaccination by BNT162b2 or CoronaVac did not improve S-specific CD4 T cell responses among 2×CorV vaccinees, positive correlations among S-specific CD4 T cells, S-specific B cells and NAbs were limited to the 3×BNT group (Figure 4A, red rectangle). However, positive correlations between S-specific cTFH cells and NAbs were observed in 3×BNT and 2×CorV+1×BNT, but not in 3×CorV (Figure 4A-C, yellow rectangles). Interestingly, in the 3×BNT group, Omicron S-specific CD4 T cell and cTFH responses exhibited stronger correlation with S-specific B cell and the broadly NAbs than those with ancestral S-specific CD4 T cell and cTFH responses (Figure 4A, yellow rectangle). We then combined all three groups for overall analysis (Figure 4D). Strong positive correlations were consistently found in NAbs titers against all 7 viral variants (Figure 4D, purple triangle). Both age and S-specific RM B cells were negatively correlated with NAb activity (Figure 4D, purple rectangle) whereas S-specific AM B cells were positively correlated with neutralizing activity (Figure 4D, green rectangle). Moreover, the frequency of S-specific AM B cells was significantly lower in infected vaccinees than uninfected vaccinees before vaccine-breakthrough infection (Figure 4E) whereas the anti-BA.2 NAb titer did not achieve statistical significance (Figure 4F). Notably, few vaccinees (2/12, 16.7%) with NAb titer higher than 1:320 became infected. We further analyzed the relationships between immune responses and clinical characteristics among our study subjects who were subsequently infected by BA.2 (Figure 4G). NAb titer was negatively correlated with hospitalization rate (Figure 4G, purple rectangle), indicating the importance of NAb in reducing COVID-19 severity. Age was positively correlated with viral negative conversion time, suggesting a longer viral clearance time among older patients (Figure 4G, black square). Notably, IFN-γ^+^CD4 T cells were negatively associated with age and viral negative conversion time (Figure 4G, red squares). In addition, hospitalization was negatively correlated with the interval between second and third dose of vaccinations and with the interval between third dose of vaccination and symptom onset, likely suggesting the importance of the optimal timing for the third dose vaccination (Figure 4G, black rectangle). These results demonstrated that the third dose vaccination-induced NAbs and T cell response contributed to reducing risk of severe clinical outcomes after infection.

**Figure 4.**
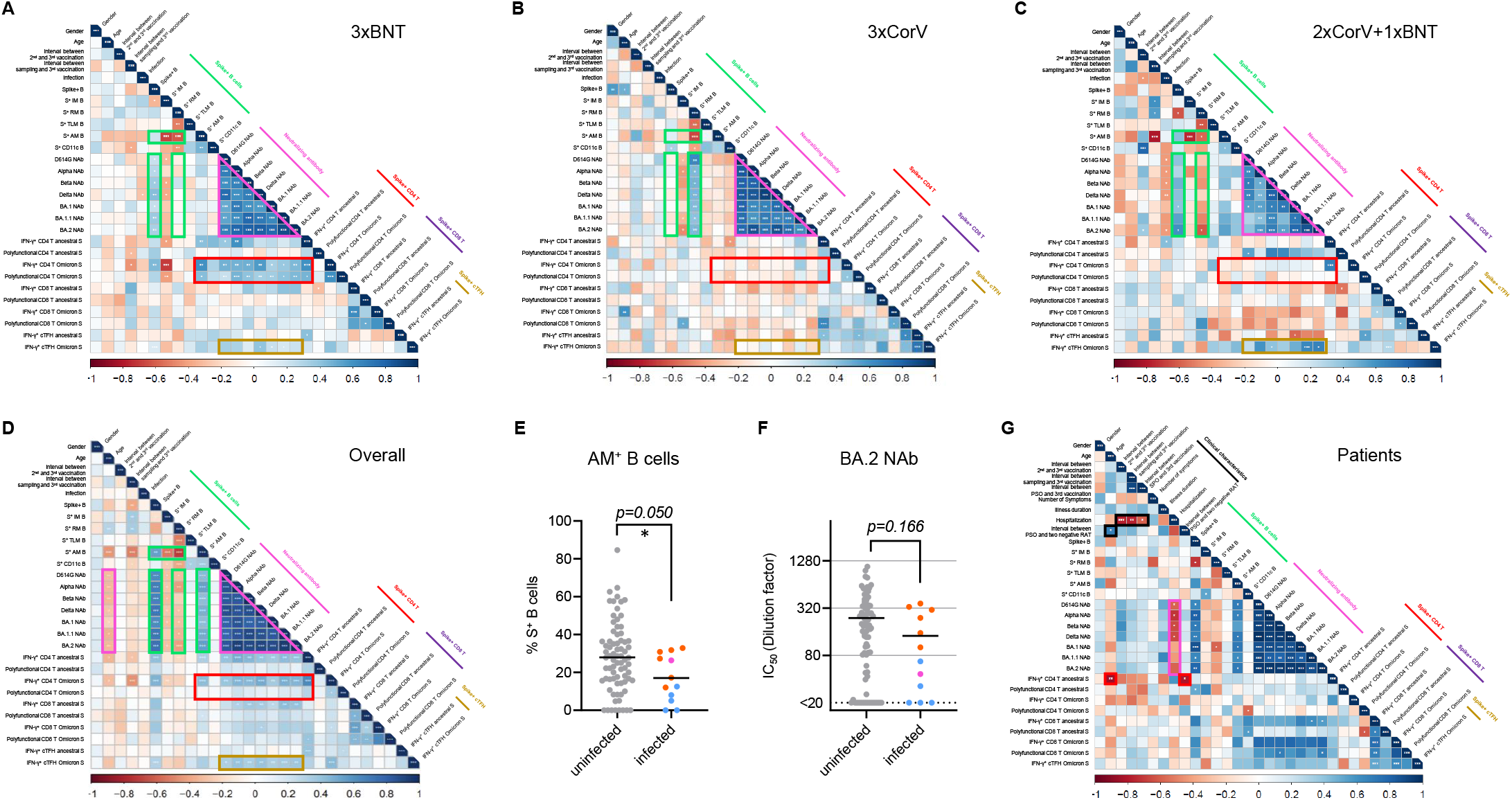
Associations among humoral, cellular immune response and breakthrough infection features. Correlogram of immune responses among 3×BNT (**A**), 3×CorV (**B**), 2×CorV+1×BNT (**C**) and overall (**D**) vaccinees. Comparison of AM^+^ B cell frequency on Spike-specific B cells (**E**) and neutralizing titer against BA.2 (**F**) between uninfected and infected vaccinees. Uninfected vaccinees, infected 3×BNT vaccinees, infected 3×CorV vaccinees and infected 2×CorV+1×BNT vaccinees were presented as grey, orange, blue and purple dots, respectively. Statistics were generated by using 2-tailed Student’s t test. *p <0.05. (**G**) Correlogram of clinical characteristics and immune responses among patients. Spearman rank order correlation values (r) are shown from red (−1.0) to blue (1.0); r values are indicated by color and square size. p values are indicated by white asterisks. The green rectangles denote SARS-CoV-2 Spike-specific B cell features, the purple triangle and rectangles denote anti-SARS-CoV-2 variants’ neutralizing antibody features, the red rectangles denote the SARS-CoV-2 Spike-specific CD4 T cell features, the yellow rectangle denotes the SARS-CoV-2 Spike-specific cTFH features and the black rectangles denotes clinical characteristic features.

### Immune responses after Omicron BA.2 breakthrough infection and the fourth vaccination

Rapidly recalled antibody and T cell responses were observed in vaccine breakthrough infections by SARS-CoV-2 variants ^17,20–22^ At median 137 (range 122-164) days post symptom onset (Supplementary Table 5), we able to harvest the blood sample from five 3×BNT, three 3×CorV and one 2×CorV+1×BNT subject who had a BA.2 breakthrough infections. Six 3×BNT, seven 3×CorV and ten 2×CorV+1×BNT subjects who never had infection were also included. For comparison, we also included three subjects who received the fourth vaccination with BNT162b2 following three-dose CoronaVac (3×CorV+1×BNT) (Supplementary Table 5). We first measured the frequency of S-specific B cells and found that BA.2 S-specific B cells were consistently lower than ancestral S-specific B cells among all vaccinees no matter with or without BA.2 infection (2.2-3.1-fold and 1.1-2.3-fold difference among uninfected and infected vaccinees, respectively) (Figure 5A-C). Among uninfected vaccinees, the frequency of BA.2 S-specific B cells in 3×CorV group (mean 0.05%) was significantly lower than those in 3×BNT (mean 0.38%) and 2×CorV+1×BNT (mean 0.17%) groups (Figure 5B). Although BA.2 infection increased BA.2 S-specific B cells in 3×CorV (mean 0.18%), it was still significantly lower than those in 3×BNT group (mean 0.53%) and lower than 3×CorV+1×BNT group (mean 0.48%) without significance (Figure 5C). In contrast to B cell response, all vaccinees showed similar CD4 and CD8 T cell responses to ancestral and Omicron Spike, and BA.2 infection did not boost a higher T cell response than uninfected vaccinees (Figure 5DI). Moreover, uninfected and infected 3×CorV showed lower T cell responses than those in 3×BNT and 3×CorV+1×BNT without significance (Figure 5F and 5I). Particularly, markedly higher CD8 T cells were found in 3×BNT uninfected vaccinees than those in 3×CorV and 2×CorV+1×BNT uninfected vaccinees even at a long term after vaccination (>4 months) (Figure 5H). These results indicated that BA.2 infection boosted cross-reactive B cells rather than T cells to ancestral and Omicron Spike.

**Figure 5.**
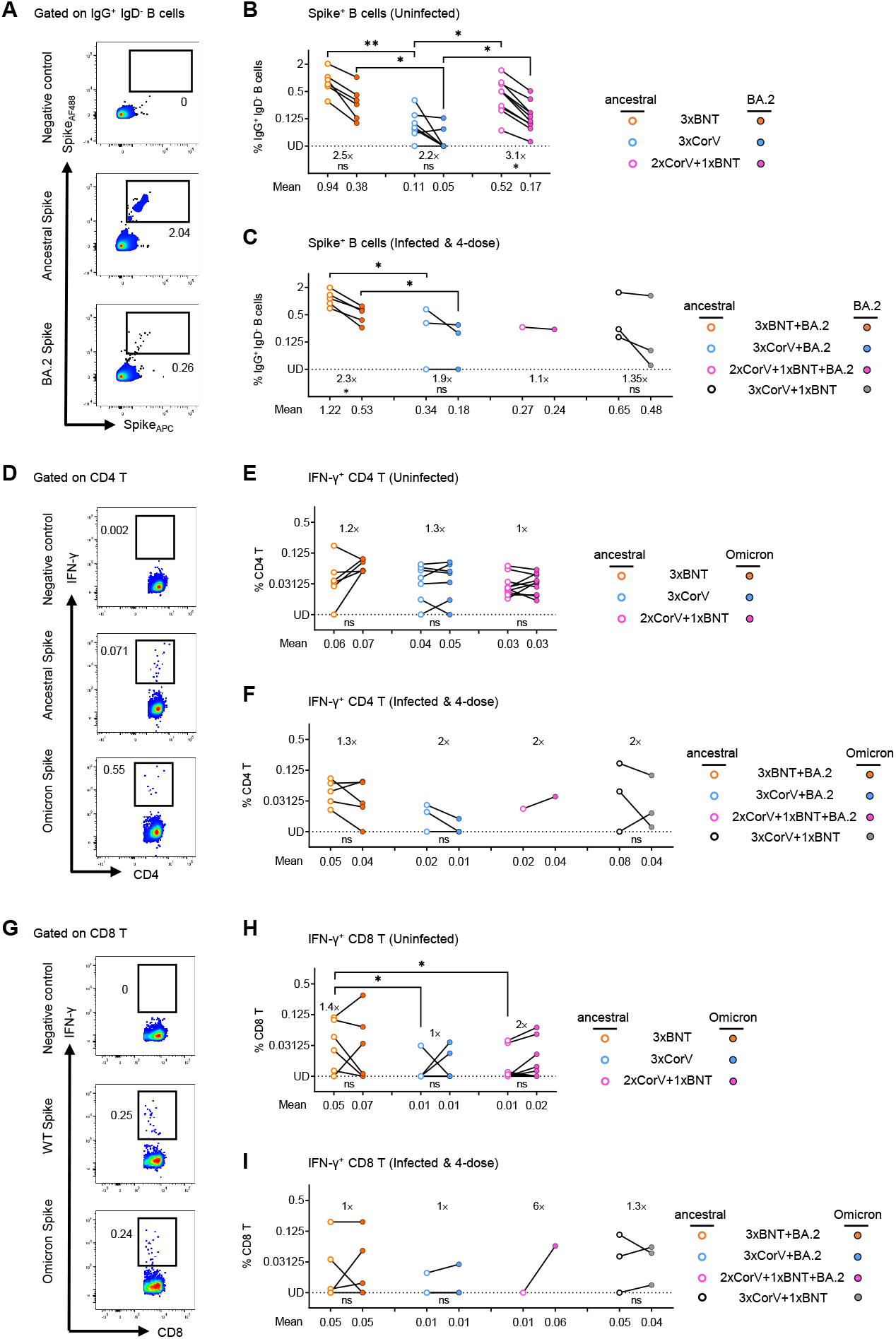
Immune responses after Omicron BA.2 breakthrough infection and the fourth vaccination. (**A**) Representative flow cytometry plots showing staining patterns of SARS-CoV-2 ancestral or BA.2 Spike probes on memory B cells (IgD^-^ IgG^+^ CD19^+^). (**B-C**) Quantified results depict the percentage of ancestral (empty) and BA.2 (solid) Spike^+^B cells in uninfected (**B**) and infected (**C**) 3×BNT (orange), 3×CorV (blue), 2×CorV+1×BNT (purple) and 3×CorV+1×BNT (grey) groups. The numbers above the x-axis indicate the fold-change in frequency of positive B cells to ancestral and BA.2 Spike. The numbers under x-axis indicate the mean frequencies of ancestral or BA.2-specific B cells. Undetected (UD): % of Spike^+^ cells<0.03125%. (**D and G**) Representative flow cytometry plots showing the IFN-γ^+^ cells among CD4^+^ (**D**) and CD8^+^ (**G**) T cells to negative control, ancestral Spike and Omicron Spike peptide pools. Quantified results depict the percentage of ancestral (empty) and Omicron (solid)-specific IFN-γ^+^ cells in uninfected (**E and H**) and infected (**F and I**) 3×BNT (orange), 3×CorV (blue), 2×CorV+1×BNT (purple) and 3×CorV+1×BNT (grey) groups. The numbers above the figures indicate the fold-change in frequency of positive T cells to ancestral and BA.2 Spike. The numbers under x-axis indicate the mean frequencies of ancestral or Omicron-specific IFN-γ^+^ cells T cells. Undetected (UD): % of IFN-γ^+^ cells<0.00781%. Each symbol represents an individual donor. Statistics were generated by using 2-tailed Student’s t test. Ns: no significance, * p<0.05; **p<0.01.

### Neutralizing antibody titer after BA.2 breakthrough infection and the fourth vaccination

Since broadly neutralizing activity would be boosted by an increased number of exposures to SARS-CoV-2 antigens (vaccination or infection) among vaccinees ^17,21,23,24^, pairwise comparison of neutralizing activity was analyzed using the plasma sample collected before (1^st^) and after (2^nd^) BA.2 breakthrough infection. Three-dose and 4-dose uninfected vaccinees were also included (Supplementary Table 5). Consistent to our previous findings in two-dose vaccinees ^7^, NAb titer of uninfected vaccinees waned over time, especially when against BA.2.12.1 and BA. 4/5 (Figure 6A-E), but the waning effect was not observed in NAbs against D614G (Figure 6A). However, 100% and 90% of the uninfected 3×BNT and 2×CorV+1×BNT vaccinees were maintained measurable NAbs against all Omicron variants whereas more uninfected 3×CorV vaccinees (4/7) loosed neutralizing capacity against Omicron BA.4/5. Notably, the fourth vaccination can boost higher NAbs titers and responder rates for 3×CorV vaccinees (Figure 6A-E). Moreover, different 3-dose vaccinees after BA.2 breakthrough infection and 3×CorV+1×BNT vaccinees consistently exhibited a stronger GMT against BA.1 (3×BNT: 3653, 3×CorV: 582 and 2×CorV+1×BNT: 221) and BA.2 (3×BNT: 3005, 3×CorV: 742 and 2×CorV+1×BNT: 417) than those against BA.2.12.1 (3×BNT: 1857, 3×CorV: 531 and 2×CorV+1×BNT: 135) and BA.4/5 (3×BNT: 957, 3×CorV: 200 and 2×CorV+1×BNT: 94) (Figure 6A-E). This boost effect by BA.2 breakthrough infection was more profound in 3×CorV vaccinees with the highest fold-change (up to 21.2-fold increased for BA.2) in GMT against Omicron sublineages (Figure 6A-E). The results indicated that BA.2 breakthrough infection and the fourth vaccination enhanced cross-neutralizing antibodies to Omicron sublineages in all vaccinees.

**Figure 6.**
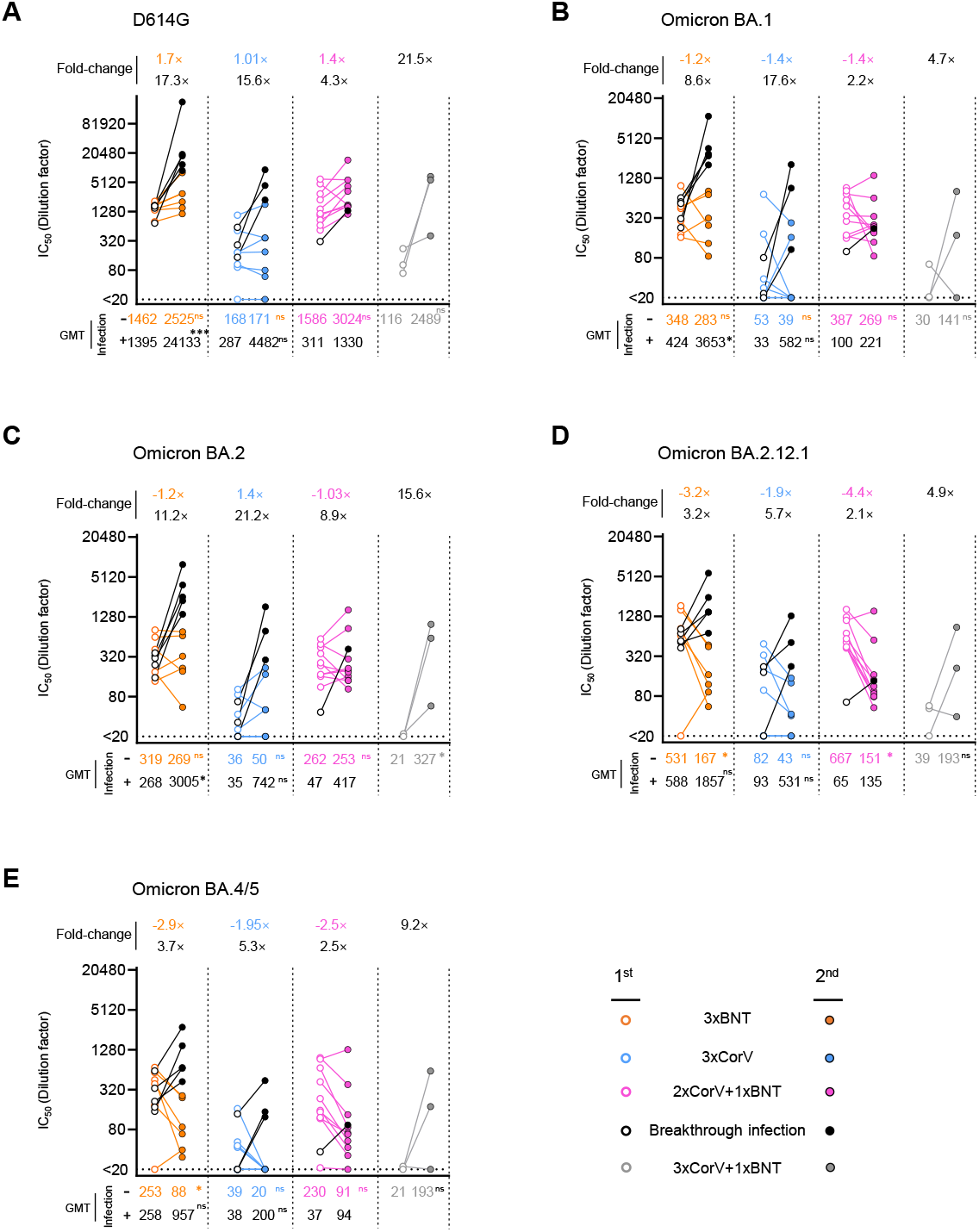
Neutralizing antibody titer after BA.2 breakthrough infection and the fourth vaccination. **(A-E)** The neutralizing antibody (IC50 represents serum dilution required to achieve 50% virus neutralization) against five SARS-CoV-2 strains were measured by pseudovirus-based assay among uninfected and infected 3×BNT (orange), 3×CorV (blue), 2×CorV+1×BNT (purple) and 3×CorV+1×BNT (grey) before (1^st^, empty dots) and after (2^nd^, solid dots) BA.2 infection or the fourth vaccination. Black dots and lines represent the breakthrough infection sample in each group. Numbers on the figure top indicate the fold-change in NAb titer between 1^st^and 2^nd^ sample. Numbers under the x-axis indicate the geometric mean titers (GMT). Statistics were generated by using 2-tailed Student’s t test. *p<0.05; ***p<0.001; ns: not significant. (**F-H**) The ratio of SARS-CoV-2 VOC NAb IC50 normalized against the D614G NAb IC50. Orange line, blue line and purple line represent uninfected 3×BNT, 3×CorV and 2×CorV+1×BNT vaccinees. Black lines represent the infected vaccinees in each group. Numbers on the figure top indicate the ratio for corresponding VOCs.

## Discussion

Clinical trials have demonstrated that a third heterologous booster vaccination by EUA SARS-CoV-2 mRNA vaccines (BNT162b2 and mRNA-1273) increased neutralizing antibody titer accompanied by better prevention and lower disease severity than the initial two doses with BBIP-CorV or CoronaVac during the Gamma and Delta epidemics^25–29^. After the emergence of the Omicron variants, some cohort studies reported that Omicron BA.1 infection was associated with milder disease and shorter duration of clinical symptoms than Delta infection^30–35^.

The third vaccination was helpful in reducing the infection and hospitalization rates during the Delta and Omicron BA.1 prevalence in other countries ^25,36,37^ Till now, the association between immune responses induced by the third vaccination and Omicron BA.2 breakthrough infection remains unknown. In this study, we investigated the immune responses of vaccinees after they received the third vaccination right before the explosive fifth wave of SARS-CoV-2 epidemic caused by Omicron BA.2 in Hong Kong ^10,14^ We also followed up the infection status and clinical outcomes of our study subjects during the wave period. We found that the third dose of either BNT162b2 or CoronaVac led to significantly lower infection rates than those who received the standard 2-dose vaccination regimen, particularly in the heterologous 2×CorV+1×BNT group. Furthermore, the third BNT162b2 resulted in significantly higher rates of asymptomatic and lower rates of hospitalization than 3×CorV group. Our findings, therefore, provided critical knowledge on understanding the role of third vaccination-induced immune responses in protection against the globally spreading Omicron BA.2 infections.

Omicron BA.2 has higher transmissibility and immune evasion than Omicron BA.1 ^38,39^, explaining its rapid spread in Hong Kong and other places ^40,41^. Since the end of January 2022, BA.2 has quickly dominated the fifth wave of SARS-CoV-2 epidemic in Hong Kong, where the universal masking policy remains unchanged, with a shorter doubling time of 1.28 days than 1.6-2.8 days of BA.1 ^10^. BA.2 shares 21 mutations in the Spike with BA.1. Although Q496S and N501Y mutations are missing in the BA.2 S-BRD domain, unique S371F, T376A, D405N and R408S mutations have been found ^39^ Due to these mutations, we and others ^39,42^ demonstrated that NAb titers against BA.2 showed 0.97-1.18 and 1.14-1.42 time lower than those against BA.1 at 0-4 weeks and >4 weeks after third vaccination by BNT162b2 or CoronaVac. Also, we consistently found that BA.2 confers the highest NAb resistance compared with other VOCs including BA.1 and BA.1.1 before emergence of BA.4/5. While 59-71% and 29-41% BNT162b2 booster recipients had low (IC50: 20-320) and median (IC50: 321-1280) NAbs against BA.2, 66% CoronaVac booster recipients had undetectable (IC50<20) NAbs. Surprisingly, although the third BNT162b2 vaccination boosted higher anti-BA.2 NAb titer and responder rate as well as a more S-specific T cell responses than the third CoronaVac, there was no significant difference in incidence of breakthrough infections between 3×BNT and 3×CorV. Firstly, the majority of our vaccinees, including 3×BNT and 3×CorV, have a low neutralizing antibody titer at the time of exposure, rendering them susceptible to BA.2 breakthrough infection. Ten of twelve vaccinees who had IC_50_<320 NAb against BA.2 became infected, which is consistent to the animal study that the prophylactic administration of convalescent plasma at 1:320 dilution hardly prevents SARS-CoV-2 infection in hamster model ^16^. Secondly, both CoronaVac and BNT162b2 hardly induce enough mucosal neutralizing antibody or T cell responses for prevention ^43^, as Omicron replicates faster and stronger than wild type and Delta variant in the nasal and bronchial compartments but less efficiently in the lung parenchyma^44–46^. Critically, although CoronaVac displays lower immunogenicity than BNT162b2, it still induced memory B cell and T cell responses that can be recalled for protection as demonstrated in the 3×CorV vaccinees with BA.2 breakthrough infection. Therefore, the recalled immune response, especially the comparable T cell responses, which are invoked by the BA.2 breakthrough infection in participants who received different vaccine regimens.

In addition, three doses of either CoronaVac or BNT162b2 vaccines provided similar and high protection against Omicron infection-induced severe outcomes ^47,48^.Such BA.2 infection-mediated immune activation might be even more profound among 3×CorV vaccinees, resulting in significantly reduced infection and hospitalization rates compared with 2×CorV vaccinees. Therefore, when all vaccinees were analyzed together, we found that S-specific activated memory B cell subset was a significant factor in preventing BA.2 infection because these AM B cells could differentiate into long-lived plasma cells ^49^ and are associated with expansion of memory B cells, and the re-establishment of B cell memory after the third vaccination ^23,50^. Moreover, T cell responses could be another protective factor because they may recognize mutated viral variants without significantly reducing the potency ^51^. We found that both BNT162b2 and CoronaVac-induced T cell responses cross-reacted to Omicron S peptides with comparable activities to ancestral S ^52,53^. Since S-specific T cells are associated with the control and clearance of the ongoing infection ^12^, potent T cell responses correlated with fewer hospitalization among patients who received the third vaccination.

While we studied the BA.2 variant, the BA.2.12.1, BA.4, and BA.5 have raised and increased resistance compared to previous VOCs to vaccine-induced NAbs through the L452R/Q and F486V mutations in the Spike^54–56^ We confirmed that BA.2 breakthrough infection and the fourth vaccination effectively boosts neutralizing antibody against BA.2.12.1 and BA.4/5. This can explain why BA.1/BA.2 infection in vaccinated persons were less at risk of BA.4/5 infection than individuals infected with a pre-Omicron VOCs ^57^. However, BA.2 breakthrough infections mainly recalled vaccine-induced ancestral Spike-specific memory B cells, which may drive further mutation of virus and variant-associated reinfection ^55,58,59,60^.

There are some limitations in our study. Firstly, most of our infected vaccinees were confirmed to have been infected by self-RAT, thus the effect of different vaccine regimens on controlling viral loads could not be determined. It remains necessary to compare the dynamics and magnitudes of the recalled immune responses among vaccinees with BA.2 breakthrough infection patients in the future. Secondly, it should be noted that the median interval time between the latest vaccination and symptom onset for the 2×BNT (227 days) and 2×CorV (237 days) groups was significantly longer than those for 3 dose vaccination groups, including 3×BNT (48.5 days), 3×CorV (56 days) and 2×CorV+1×BNT (25.5 days). Although NAb potency wans over time ^7^, we and others consistently found that timely boost vaccination not only restore waning NAb titers but also broaden the breadth of NAbs, which is able to cross-neutralize VOCs, including Omicron ^8,23,50,61^. Thirdly, only one sample can be harvested from 2×CorV+1×BNT vaccinees with BA.2 infections. It’s hard to conclude the outcome of BNT162b2 booster for two-dose CoronaVac vaccinees during BA.2 breakthrough infection.

In summary, we report that 3×BNT and 3×CorV provided better protection against SARS-COV-2 BA.2 than 2×BNT and 2×CorV. High frequencies of S-specific activated memory B cells and cross-reactive T cell responses induced by the third vaccination are critical for protection and illness reduction during the Omicron BA.2 breakthrough infection. Enhanced neutralization induced by BA.2 breakthrough infection and the fourth vaccination may help to reduce the risk for infection of ongoing BA.2.12.1 and BA.4/5.

## Contributors

Z.C. and R.Z. conceived and designed the study. R.Z. and Z.C. designed experiments, analyzed data, and wrote the manuscript. Z.C., R.Z. X.L., Y-C.K., H.-Y.K., I.F.-N.H., and K.-Y.Y coordinated donor recruitment and specimen collection. R.Z., N.L., H.H., D.Y., Q.P. prepared the clinical sample. R.Z., N.L. and H.H. performed the flow cytometry analysis. R.Z., N.L. and D.Y performed the pseudoviral neutralization assay. Z.D. did the correlation analysis.

## Data sharing

The authors declare that the data supporting the findings of this study are available from the corresponding author upon request.

## Acknowledgements

We sincerely thank Drs. David D. Ho and Pengfei Wang for the expression plasmids encoding for D614G, Alpha, Beta variants and Omicron sub-lineages, Dr. Linqi Zhang for the Delta variant plasmid, Mrs. Tsz-Tat Chan and Mark Wai-Kwan Woo for helping with the survey. We thank Mr. Shek-Hong Kwan for editorial work. This study was supported by the Hong Kong Research Grants Council Collaborative Research Fund (C7156-20G, C1134-20G and C5110-20G) and Health and Medical Research Fund (19181012, COVID1903010 and COVID190123); Wellcome Trust (P86433); Shenzhen Science and Technology Program (JSGG20200225151410198 and JCYJ20210324131610027); the Health@InnoHK, Innovation and Technology Commission of Hong Kong and the China National Program on Key Research Project (2020YFC0860600, 2020YFA0707500 and 2020YFA0707504); Emergency Key Program of Guangzhou Laboratory (EKPG22-01) and donations from the Friends of Hope Education Fund. Z.C.’s team was also partly supported by the Hong Kong Theme-Based Research Scheme (T11-706/18-N).

## Supplementary materials

**Supplementary Table 1.** Significance in demographic characteristics among each vaccine cohort.

|  | P value |  |  |  |  |  |  |
| --- | --- | --- | --- | --- | --- | --- | --- |
|  | 2×BNT<br>vs<br>3×BNT | 2×CorV<br>vs<br>3×CorV | 2×CorV<br>vs<br>2×BNT | 3×CorV<br>vs<br>3×BNT | 2×CorV<br>+1×BNT<br>vs<br>2×CorV | 2×CorV<br>+1×BNT<br>vs<br>3×CorV | 2×CorV<br>+1×BNT<br>vs<br>3×BNT |
| Vaccinations |  |  |  |  |  |  |  |
| Infection rate % | <0.0001 | 0.009 | 0.828 | 0.22 | <0.0001 | 0.159 | 0.426 |
| Age | <0.0001 | 0.2654 | 0.0006 | 0.1024 | 0.6595 | 0.9274 | 0.4714 |
| Gender | 0.21 | 1.0 | 0.294 | 1.0 | 0.515 | 0.524 | 1.0 |
| Median interval days<br>between latest<br>vaccination and symptom<br>onset | <0.0001 | <0.0001 | 0.4355 | 0.3926 | 0.0027 | 0.1598 | 0.2162 |
| Asymptomatic rate % | 0.821 | ns | 1.0 | ns | ns | ns | ns |
| Number of symptoms | 0.3009 | 0.2634 | 0.7435 | 0.6130 | 0.8218 | 0.3983 | 0.5026 |
| Presentation to hospital % | 0.183 | 1.0 | 1.0 | 0.541 | 0.426 | 0.371 | 0.163 |
| Duration of illness, days | 0.8024 | 0.4392 | 0.1780 | 0.8306 | 0.9264 | 0.6639 | 0.8108 |
| The interval days between<br>symptom onset and two<br>negative RAT | 0.9501 | 0.474 | 0.3277 | 0.8324 | 0.9645 | 0.7541 | 0.5315 |

**Supplementary Table 2.** Characteristics of the third doses of SARS-CoV-2 vaccinee cohorts.

| Characteristics | 2xBNT<br>(n=27) | 2xCorV<br>(n=16) | 3xBNT<br>(n=41) | 3xCorV<br>(n=28) | 2xCorV+1xBNT<br>(n=21) |
| --- | --- | --- | --- | --- | --- |
| Age | 30 (22-66) | 27 (22-33) | 46 (27-55) | 51 (40-58) | 47 (32-53) |
| Gender |  |  |  |  |  |
| Male (n) | 11 | 10 | 29 | 15 | 19 |
| Female (n) | 16 | 6 | 12 | 13 | 2 |
| Days between the 1st and 2nd dose | 21 (20-31) | 28 (22-35) | 23 (21-36) | 28 (28-71) | 29 (28-97) |
| Days between the 2nd and 3rd dose | - | - | 236<br>(180-283) | 236 (191-287) | 240 (189-284) |
| Days between last vaccination and blood collection | 31 (7-47) | 27 (10-105) | 23 (7-75) | 56 (13-77) | 47 (7-77) |
| Number of Infection after last vaccination | 0 | 0 | 6 | 5 | 1 |
| Values displayed are medians, with ranges in parentheses |  |  |  |  |  |

**Supplementary Table 3.**
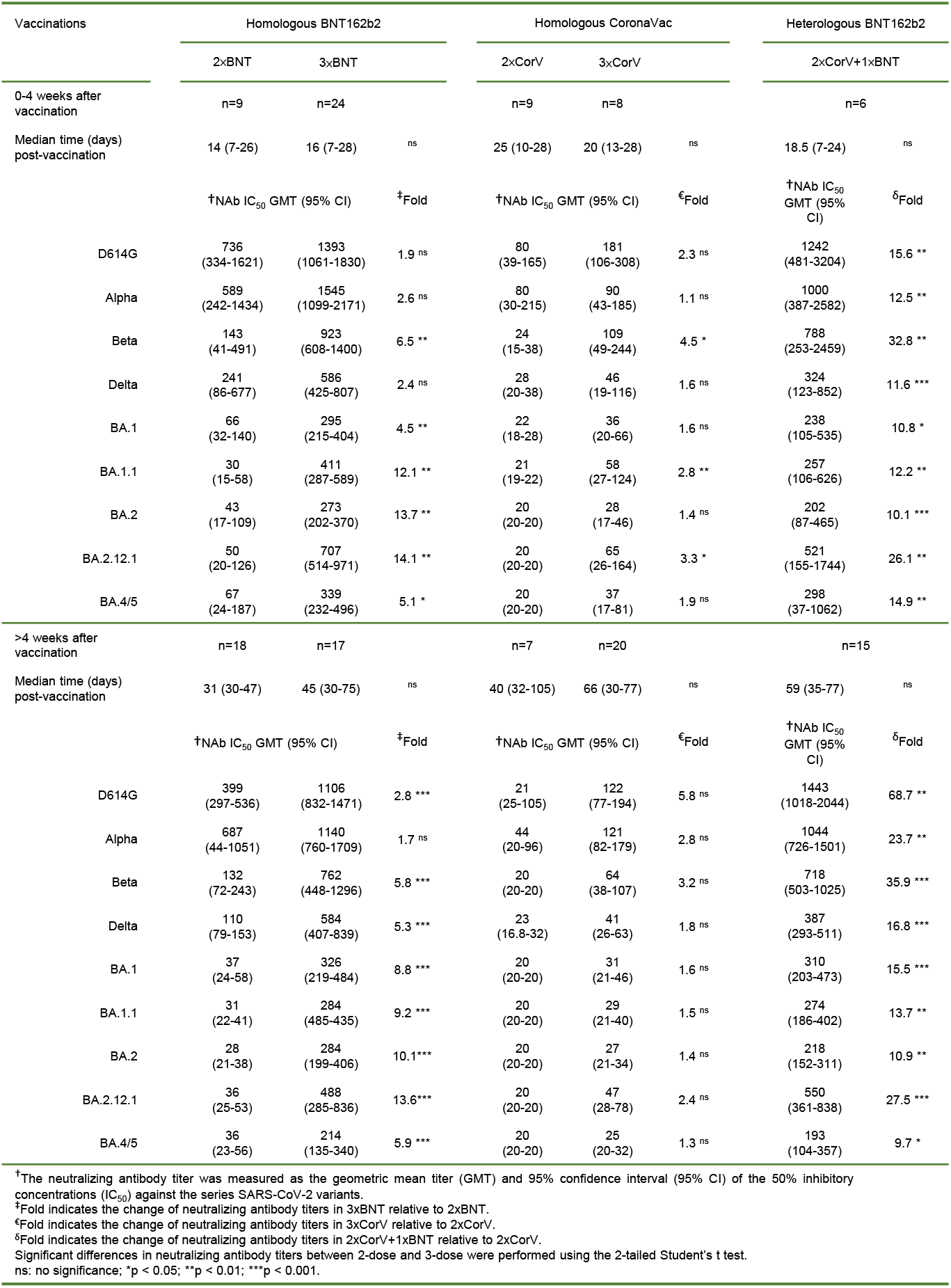
Comparison in neutralizing antibody titers between 2-dose and 3-dose vaccinations.

**Supplementary Table 4.** Comparison in antibody responder rates between 2-dose and 3-dose vaccination.

| Vaccinations | Homologous BNT162b2 |  | Homologous CoronaVac |  | Heterologous BNT162b2 |
| --- | --- | --- | --- | --- | --- |
|  | 2xBNT | 3xBNT | 2xCorV | 3xCorV | 2xCorV+1xBNT |
| 0-4 weeks after vaccination |  |  |  |  |  |
| Median time (days) post-vaccination | 14 (7-26) | 16 (7-28) | 25 (10-28) | 20 (13-28) | 18.5 (7-24) |
| Responder rate % (No. participants with response/Total No.) |  |  |  |  |  |
| D614G | 100<br>(9/9) | 100<br>(24/24) | 89<br>(8/9) | 100<br>(8/8) | 100<br>(6/6) |
| Alpha | 100<br>(9/9) | 100<br>(24/24) | 67<br>(6/9) | 100<br>(8/8) | 100<br>(6/6) |
| Beta | 67<br>(6/9) | 100<br>(24/24) | 11<br>(1/9) | 88<br>(7/8) | 100<br>(6/6) |
| Delta | 89<br>(8/9) | 100<br>(24/24) | 56<br>(5/9) | 63<br>(5/8) | 100<br>(6/6) |
| BA.1 | 67<br>(6/9) | 100<br>(24/24) | 11<br>(1/9) | 88<br>(7/8) | 100<br>(6/6) |
| BA.1.1 | 22<br>(2/9) | 100<br>(24/24) | 11<br>(1/9) | 63<br>(5/8) | 100<br>(6/6) |
| BA.2 | 33<br>(3/9) | 100<br>(24/24) | 0<br>(0/9) | 38<br>(3/8) | 100<br>(6/6) |
| BA.2.12.1 | 67<br>(6/9) | 100<br>(24/24) | 0<br>(0/9) | 63<br>(5/8) | 100<br>(6/6) |
| BA.4/5 | 56<br>(5/9) | 100<br>(24/24) | 0<br>(0/9) | 38<br>(3/8) | 100<br>(6/6) |
| >4 weeks after vaccination |  |  |  |  |  |
| Median time (days) post-vaccination | 31 (30-47) | 45 (30-75) | 40 (32-105) | 66 (30-77) | 59 (35-77) |
| Responder rate % (No. participants with response/Total No.) |  |  |  |  |  |
| D614G | 100<br>(18/18) | 100<br>(17/17) | 86<br>(6/7) | 90<br>(18/20) | 100<br>(15/15) |
| Alpha | 100<br>(18/18) | 100<br>(17/17) | 71<br>(5/7) | 100<br>(20/20) | 100<br>(15/15) |
| Beta | 89<br>(16/18) | 100<br>(17/17) | 0<br>(0/7) | 70<br>(14/20) | 100<br>(15/15) |
| Delta | 94<br>(17/18) | 100<br>(17/17) | 71<br>(5/7) | 50<br>(10/20) | 100<br>(15/15) |
| BA.1 | 50<br>(9/18) | 100<br>(17/17) | 0<br>(0/7) | 50<br>(10/20) | 100<br>(15/15) |
| BA.1.1 | 39<br>(7/18) | 100<br>(17/17) | 0<br>(0/7) | 30<br>(6/20) | 100<br>(15/15) |
| BA.2 | 39<br>(7/18) | 100<br>(17/17) | 0<br>(0/7) | 35<br>(7/20) | 100<br>(15/15) |
| BA.2.12.1 | 56<br>(10/18) | 94<br>(16/17) | 0<br>(0/7) | 50<br>(10/20) | 100<br>(15/15) |
| BA.4/5 | 33<br>(6/18) | 94<br>(16/17) | 0<br>(0/7) | 25<br>(5/20) | 93<br>(14/15) |

**Supplementary Table 5.**
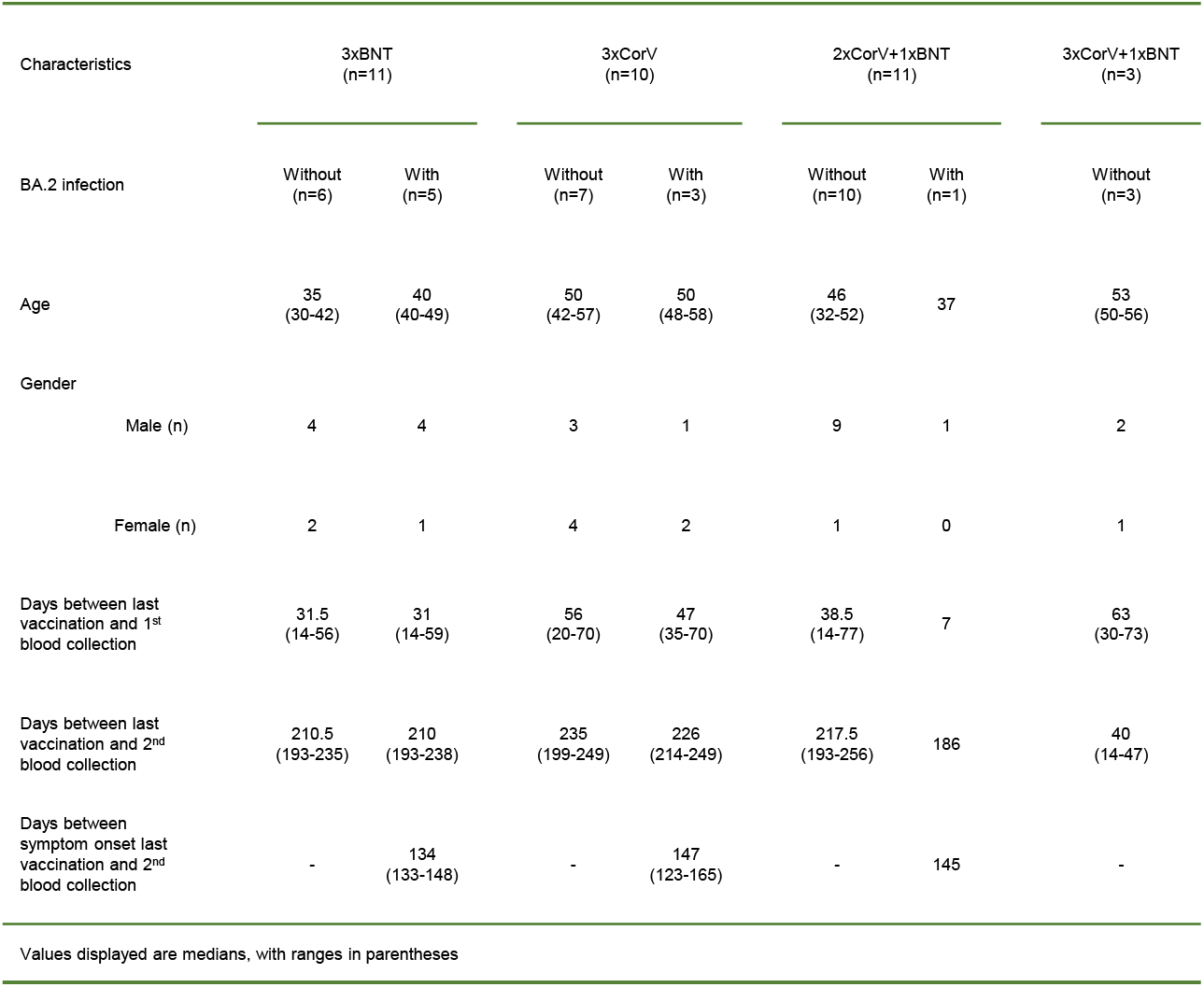
Characteristics of three doses and 4 doses of SARS-CoV-2 vaccinees with or without BA.2 infection.

**Supplementary Figure 1.**
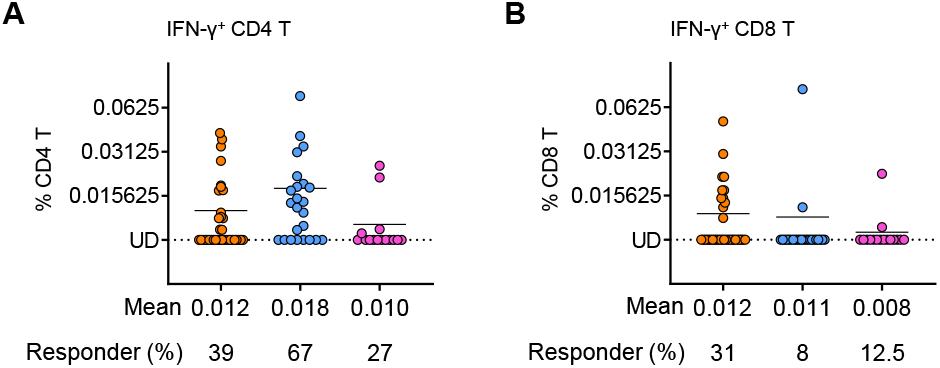
SARS-CoV-2 NP-specific T cell responses. PBMCs from vaccinees were subjected to the intracellular cytokine staining assay against NP peptide pool. IFN-γ^+^ cells were gated on CD4 (**A**) and CD8 (**B**) T cells, respectively. Quantified results depict the percentage of IFN-γ^+^ cells as background subtracted data from the same sample stimulated with negative control (anti-CD28/CD49d only). Each symbol represents an individual donor with a line indicating the mean of each group among the 3×BNT (orange), 3×CorV (blue) and 2×CorV+1×BNT (purple) vaccinees. The mean frequency of IFN-γ^+^ cells and responder rates were depicted under x-axis (% of IFN-γ+ cells>0.00781% termed ‘responder’ after subtracted from percentage of unstimulated control). Undetected (UD): % of IFN-γ^+^ cells<0.00781%.

**Supplementary Figure 2.**
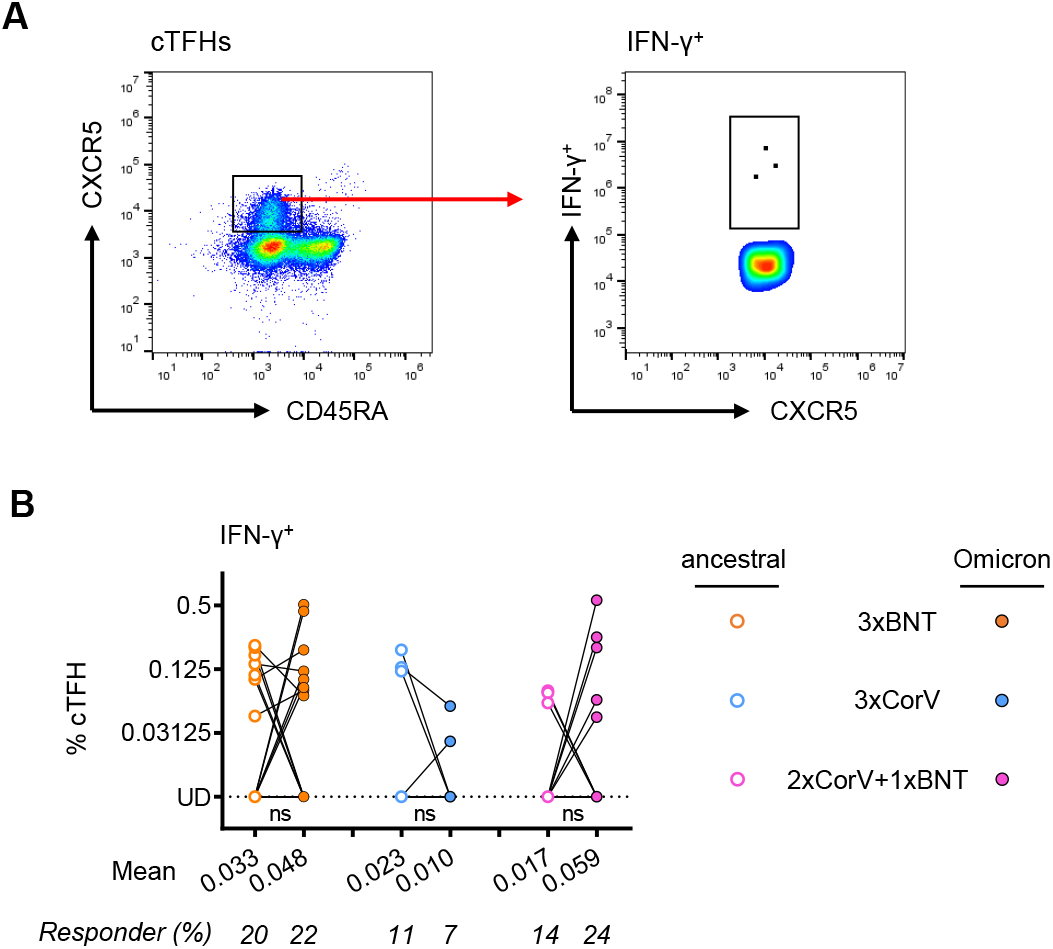
SARS-CoV-2 Spike-specific cTFH responses. PBMCs from vaccinees were subjected to the intracellular cytokine staining assay against Spike peptide pools from ancestral or Omicron SARS-CoV-2. (**A**) IFN-γ^+^ cells were gated on cTFHs. (**B**) Quantified results depict the percentage of IFN-γ^+^ cells as background subtracted data from the same sample stimulated with negative control (anti-CD28/CD49d only). Each symbol represents an individual donor with a line indicating the mean of each group to ancestral (open dots) or Omicron (solid dots) Spike among the 3×BNT (orange), 3×CorV (blue) and 2×CorV+1×BNT (purple) vaccinees. The mean frequency of IFN-γ^+^ cells and responder rates were depicted under x-axis. Undetected (UD): % of IFN-γ^+^ cells<0.00781%. Statistics were generated by using 2-tailed Student’s t test. Ns: no significanceNs: no significance.

